# Metabolic cost sets the shear stress profile of optimal vascular trees

**DOI:** 10.64898/2026.09.29.755319

**Authors:** Genaro Vázquez-Victorio, Enrique Hernández-Lemus, Alejandra Cervera, Laura Gómez-Romero, Hugo Tovar, Daniel Pérez-Calixto

**Affiliations:** Facultad de Ciencias, Universidad Nacional Autónoma de México, Mexico City, Mexico; Instituto Nacional de Medicina Genómica, Mexico City, Mexico

## Abstract

Vascular remodeling responds to local wall shear stress, but transport efficiency belongs to the network as a whole. We connect these scales by minimizing dissipation under a fixed maintenance budget, with vessel cost proportional to mass raised to an exponent *b*. At an optimum, dissipated power is proportional to vessel cost. This yields the shear set-point | *τ* | ∝ *r*^*b*−1^*l*^(*b*−1)/2^ for radius *r* and length *l*, with uniform shear recovered at *b* = 1. The associated local adaptation rule decreases the global functional, although the topology it reaches retains a memory of development. Simulations with 400 and 700 sinks show that staged growth improves transport mainly by allowing pruned connections to regrow. Fluctuating demand sustains loops without increasing the maintenance budget. The local condition fixes allocation on a given support, whereas growth and variable demand shape the topology that survives.

---

A basic question runs through the study of adaptive transport networks: how can local remodeling rules achieve a global energetic goal? The question arises in vascular systems and leaf venation, but also in other biological flow networks and engineered transport systems. Blood vessels make the tension especially clear. They sense flow-induced stress through mechanotransduction [1]; in the classic experiment of Kamiya and Togawa, increasing carotid flow caused dilation, after which wall shear stress moved back toward its initial value [2]. Shear stress, however, is not uniform across a vascular bed [3, 4]. What local target should a vessel follow when the network must balance dissipation against maintenance cost?

Murray’s principle gives one answer when maintenance cost is proportional to blood volume [5]. Optimization models extend that idea, linking vessel geometry to allometric scaling [6]. With a fixed flow pattern, concave transport costs favor trees [7–9]; fluctuating loads and damage may instead favor loops [10–12]. Here a power-law maintenance cost sets a shear target that depends on both radius and length. On any prescribed tree, the target determines the optimal allocation exactly, yet it leaves the topology open. We follow the networks selected by local adaptation to see how staged growth and fluctuating demand shape their architecture.

## Network and cost

*—*Take a connected graph with *N* sinks, a single source *s*, and *n* = *N* + 1 nodes. Each sink withdraws one unit of flow. An edge *e* has radius *r*_*e*_, length *l*_*e*_, and Poiseuille conductance. Absorbing viscosity and fixed geometric prefactors into the reference units gives 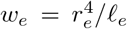 for the conductance, *f*_*e*_ for the edge flow, and 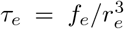 for the signed wall shear stress. None of the exponents derived below depends on this choice of units.

Let ***b***_*e*_ denote the oriented incidence vector of edge *e*, with entries +1 and −1 at its endpoints, and let ***e***_*i*_ be the unit vector at node *i*. Then 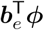 is the pressure drop across the edge. The injection vector is ***I*** = *N* ***e***_*s*_ − Σ _*t ≠ s*_ ***e***_*t*_, and the weighted graph Laplacian is 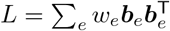. Up to a common offset, node pressures are ***ϕ*** = *L*^+^***I***, where *L*^+^ is the Moore–Penrose pseudoinverse. Thus 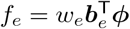 and

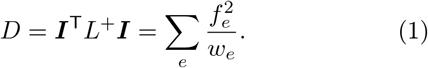

For vessel mass 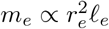, assume a maintenance cost proportional to 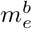. In the same reference units, the total cost is

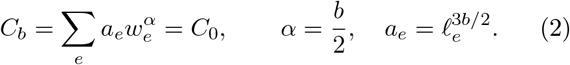

Here *C*_0_ *>* 0 is the prescribed budget and 0 *< b <* 2; *b* = 1 gives the usual volume cost. Along with the Kleiber and Rubner exponents, 3*/*4 and 2*/*3, we consider the more concave choices 1*/*2 and 1*/*4. The exponent *b* is treated as a constitutive property of vessel maintenance, motivated by metabolic scaling at the whole-organism level.

Because the cost is nonlinear in mass, segmentation matters. Splitting a segment of mass *m* into two equal parts changes its cost from *m*^*b*^ to 2^1*−b*^*m*^*b*^. Our edges run between demand nodes; an anatomical test would need a consistent rule of its own, such as defining segments between branch points.

Once the conductances are eliminated, a prescribed flow pattern leads to a concave flow cost for 0 *< b <* 2, so an optimum may be supported on a tree [7–9]. Downstream demand fixes the flows on that tree, and the optimal allocation is explicit (Appendix A). Network com-parisons use 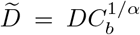, an invariant under uniform rescaling of all radii.

*The local optimality condition*.*—*Let *λ >* 0 be the multiplier for Eq. (2). On an active edge, *∂D/∂w*_*e*_ =− (Δ*p*_*e*_)^2^, where 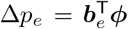 is the pressure drop. Stationarity of *D* + *λC*_*b*_ then gives

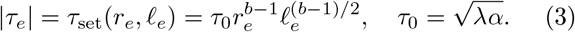

This is the central result of the optimization: the constitutive exponent of a global maintenance cost determines the local mechanical set-point. Equation (3) is necessary at an optimum on its active support. For a fixed tree it is sufficient as well, since the variables 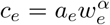 make dis-sipation strictly convex under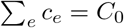. Yet the same condition can hold on different trees. The local optimum is therefore topologically degenerate.

There is also a direct energetic interpretation. Writing the power dissipated by an edge as 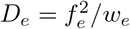 gives

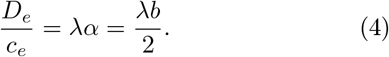

Every active vessel dissipates power in proportion to its maintenance cost. Murray’s uniform-shear condition is not universal; it is the *b* = 1 member of a broader family of optimal stress laws. When *b <* 1, both radius and length enter the set-point, and the flow law becomes 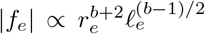. A branching exponent based only on radius therefore requires an assumption about length. At fixed length, the exponents 2.75 and 2.67 exceed the pooled coronary flow–diameter exponent of 2.39 reported in Ref. [13], although morphometry by itself cannot establish the maintenance law [14]. The shear profiles in Fig. 1 show how geometry changes the set-point across cost exponents.

**FIG. 1.**
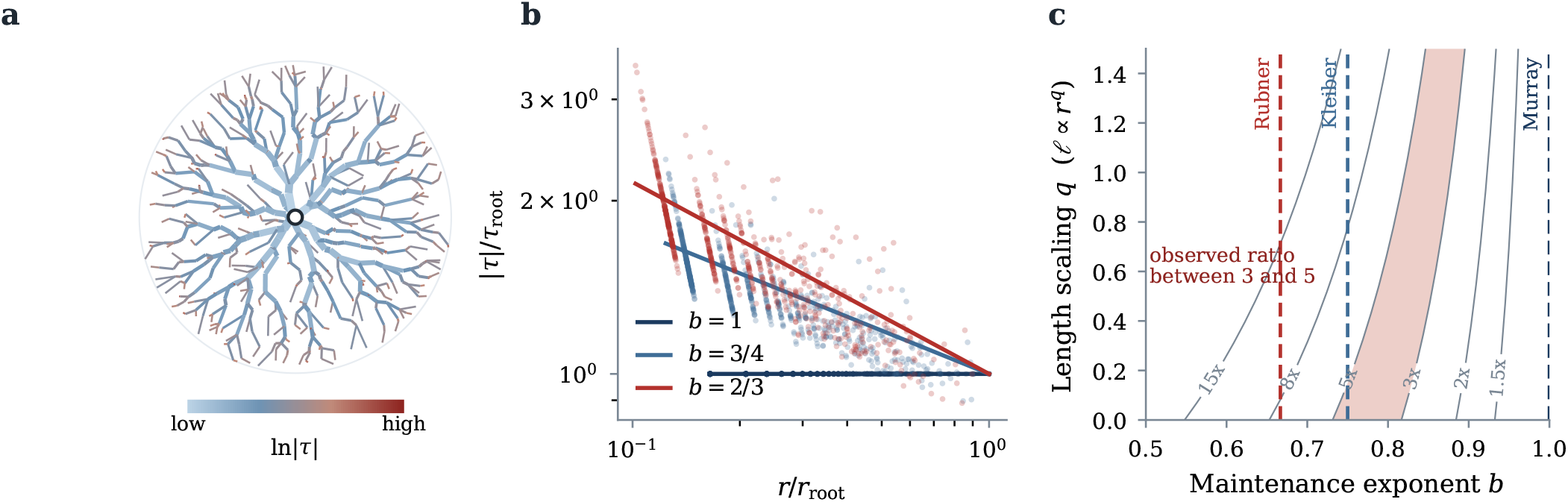
Shear set-points and the role of geometry. (a) Lowest-dissipation tree found for *N* = 700 and *b* = 3*/*4; line width denotes radius, and color denotes ln |*τ* |. (b) Shear profiles for three cost exponents. Differences in vessel length produce the spread around the radius-dependent trend. (c) Predicted ratio of shear in small and large vessels for a radius ratio *R* = 400, shown as a function of *b* and the length–radius exponent *q*. The shaded band covers ratios from 3 to 5. Panels (a) and (b) show results from the current campaign at *N* = 700.

Length cannot be ignored when comparing the theory with vascular data. A shear ratio alone does not determine *b*. If *l* ∝ *r*^*q*^ and *R* = *r*_large_*/r*_small_, then

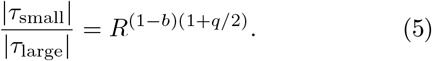

For *R* = 400 and *b* = 3*/*4, the ratio is 4.47 at *q* = 0 but about 9.5 at *q* = 1. The physiological range of roughly 3–5 discussed in Refs. [3, 4] therefore constrains cost and geometry jointly. Testing Eq. (3) requires shear stress, radius, and segment length to be measured in the same vessels (Appendix D). Its length dependence raises a further mechanobiological question: how is segment scale encoded locally?

### Local adaptation

*—*The optimality condition suggests how a vessel might adapt. Define the logarithmic radius *x*_*e*_ = ln *r*_*e*_ and the normalized shear *z*_*e*_ = | *τ*_*e*_ | */τ*_set_≥ 0, with *z*_*e*_ = 1 at the target. For a common adaptation rate *κ >* 0, consider

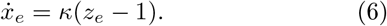

For fixed geometry, loading, active support, and *λ*, the functional *F* = *D* + *λC*_*b*_ obeys

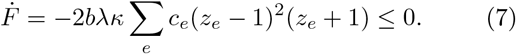

A purely local stress-sensing rule therefore performs descent on the global transport functional. Equality is reached only when every active edge is at its set-point. Hu and Cai established the broader link between local adaptation and global transport energy [15]; here, the maintenance law determines the exponents of a target written directly in terms of wall shear stress. Equation (7) holds while the support remains fixed. Pruning and growth change that support.

Does local descent also select an efficient topology? To find out, we integrate Eq. (6) in ln *r* and prune any vessel whose radius falls below 10^*−*3^ of the largest. Figure 2 follows the comparison from dense initial networks to staged growth and fluctuating demand. The campaign covers five values of *b* in three spatial domains for each size, *N* = 400 and 700. Performance is measured against the lowest-dissipation tree found, using the percentage 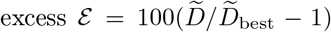. Integration parameters and convergence records appear in Appendix B.

**FIG. 2.**
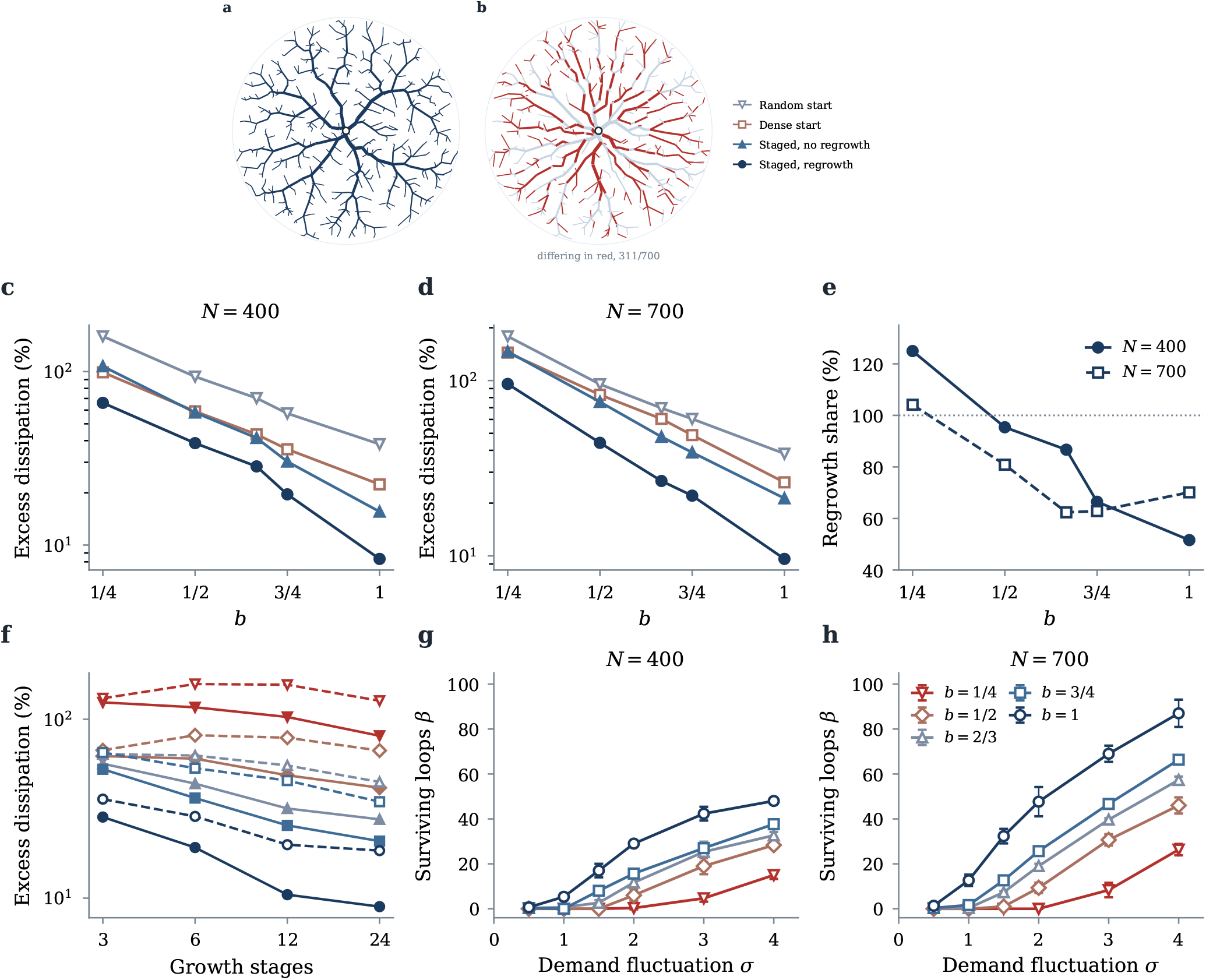
Effects of developmental history and fluctuating demand. (a) Lowest-dissipation tree found for *b* = 3*/*4 at *N* = 700; line width denotes radius. (b) Network produced by local adaptation from a dense initial state on the same domain. Edges not shared with panel (a) are shown in red, and their number is given in the panel. (c),(d) Mean excess dissipation for *N* = 400 and 700. Staged protocols use 24 growth stages and the default seeding radius. Means include three domains; random starts also include eight initializations per domain. (e) Regrowth share *S*, defined in Eq. (8). Values above 100% indicate that growth without regrowth performs worse than dense adaptation. (f) Excess dissipation versus stage count, averaged over both network sizes and three domains per size. Solid lines allow regrowth, whereas dashed lines do not. Colors in (f)–(h) identify *b*, as shown in (h). (g),(h) Mean loop count for each size, with sample standard deviations across three domains. The fixed-set-point and matched-budget protocols give identical loop counts for all 180 paired realizations; therefore, only one set of curves is shown.

Dense initial networks consistently settle on trees with higher dissipation, with a wider gap for more concave maintenance costs [Figs. 2(c) and 2(d)]. Random initial radii perform worse still (Appendix B). The resulting rest points occupy different supports: satisfying the local condition does not close the topological gap. Even when every active edge obeys the same optimality condition, network architecture retains a record of its developmental history.

### Growth and regrowth

*—*The topological gap makes development part of the problem. Growth can steer a locally adapting network toward more efficient architectures [16], and retinal angiogenesis offers a biological parallel: endothelial cells leave low-shear segments, stabilizing high-shear branches as nearby vessels regress [17]. Our rule captures this selection at the network level. Sinks are added in outward stages, with adaptation af-ter each stage; by default, previously pruned edges are also reseeded. Over 24 stages, this protocol outperforms dense adaptation for every exponent and both network sizes. The mean excess at *b* = 1 falls to 8.3% and 9.6% for *N* = 400 and 700, respectively. Even at *b* = 1*/*4, it falls to 66.1% and 95.6%. Because each mean covers three spatial domains, the spread measures variation between domains rather than confidence intervals.

To distinguish territorial expansion from the reopening of pruned connections, we repeat the protocol with regrowth disabled. Let 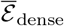, 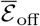, and 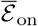denote the three-domain mean excesses for dense adaptation and for staged growth without and with regrowth. The fraction of improvement lost is

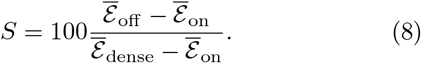

Across the ten size–exponent combinations, *S* ranges from 51.6 to 125.0% [Fig. 2(e)]. Outward growth still helps in eight combinations when regrowth is disabled. At *b* = 1*/*4, however, it raises the excess by 8.3 and 2.0 percentage points. Reopening connections accounts for much of the full protocol’s advantage.

Nor can isotropic dilation alone explain the gain; its effect varies with the exponent. A fivefold increase in the initial seeding parameter changes the 24-stage excess by only −2.45 to +0.77 percentage points, although it can reduce the three-stage excess by 22.8 points.

Regrowth is not simply an extra search step. On a fixed domain, with every sink present from the start, we first adapt to a rest point, then restore all pruned edges and adapt again. At *b* = 3*/*4 and *N* = 400, the excess falls from 39.1% to 36.1% after three rounds and to 34.3% after six, where it plateaus: twelve and twentyfour rounds also give 34.3%. Staged growth on the same domain and against the same reference reaches 17.6%, and staged growth without regrowth reaches 26.6%. The larger gain comes from reopening connections in the sequence imposed by staged construction.

The comparison survives a fourfold reduction of the integration step. At *b* = 1, the dense excess decreases from 24.6% to 18.3%, whereas the staged excess decreases from 8.5% to 7.2%. Although the absolute values shift, staged growth retains a substantial advantage (Appendix B).

### Fluctuating demand

*—*Growth decides which connections survive; variable demand can keep some of them open. Let each sink have mean demand one and independent fluctuations of variance *σ*^2^, with the source balancing the total demand. The loading second moment and expected dissipation are

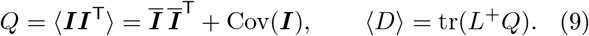

Writing the loading through *Q* links the vascular model to resistance optimization under general loading [18]. Brackets denote an average over demand, while *Q***1** = 0 expresses flow conservation. Adaptation responds to the root-mean-square shear, 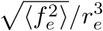. A connected support with *M*_*a*_ active edges has *β* = *M*_*a*_ − *n* + 1 independent loops. As in earlier transport models, fluctuations preserve loops [10, 11, 19]; more concave costs generally preserve fewer of them [Figs. 2(g) and 2(h)]. Environmental variability can therefore stabilize redundant topology without increasing the maintenance budget. Only a few loops survive near onset, so the present sampling cannot locate the transition precisely. Complete loop counts are given in Appendix B.

When *τ*_0_ is adjusted to match the reference budget, every paired loop count remains unchanged and all final budgets lie within 2.7 ×10^*−*6^ of one. The loops arise from fluctuating demand, not from the larger budgets reached at fixed *τ*_0_. Because budget matching begins from the adapted network, the comparison isolates material cost rather than developmental history.

## Discussion

*—*A global energetic constraint can generate an exact local mechanical rule in an adaptive transport network. With a power-law maintenance cost, both the stress set-point and the dissipation-to-cost ratio are fixed on every active edge. Topology remains outside that local rule: developmental history selects among admissible supports, whereas fluctuating demand stabilizes redundant connections.

Recent work moves beyond Murray’s universal exponent by adding vessel-wall metabolism to a single-vessel calculation [20]. Our result works at the network level. A maintenance budget shared across vessels fixes the exponent of the local shear target in Eq. (3), a prediction that calls for radius, length, and flow to be measured in the same vessels; a shear ratio alone is not enough. Viscosity, pulsatility, and the choice of vessel segmentation will matter as well. Numerically, reducing the integration step fourfold at *b* = 1 changes the absolute excesses, but not their ordering or the clear advantage of staged growth. That ordering persists across all five exponents, supporting the view that developmental history selects lower-dissipation rest points. Larger networks and finer integration will be needed to resolve asymptotic scaling and precise excess values. More broadly, the results separate three levels of organization in adaptive transport systems: energetic constraints determine local allocation, developmental history determines topology, and environmental variability determines redundancy.

This work was supported by DGAPA-PAPIIT under Grant No. IN234029 and by SECIHTI under Grant No. CBF-2025-G-789. Claude Opus 5 (Anthropic) was used as a coding assistant and OpenAI Codex for language editing; their use in the numerical implementation is described in Appendix B. The authors are responsible for the scientific content and its verification.

## Supporting information

summary.tsv

## DATA AVAILABILITY

The analysis code is available in the public repository [21]. The numerical dataset contains 1110 records and is included in the manuscript source package as summary.tsv. The quantitative panels in Figs. 1 and 2 were generated from this table. The network illustrations were recomputed with the same code and integration settings. No new experimental data were collected.

## END MATTER

### Appendix A: Allocation and numerical reference

For a fixed tree, let 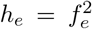 under steady demand, or 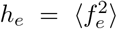 under fluctuating demand. Minimizing Σ_*e*_*h*_*e*_ */w*_*e*_ subject to Eq. (2) gives

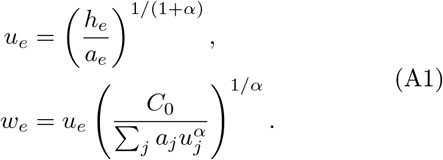

The positive quantities *u*_*e*_ set the relative conductances, while the second factor fixes the budget. On a tree with *k*_*e*_ downstream demands of unit mean and independent variance *σ*^2^,

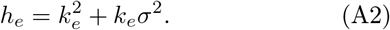

For the fluctuating reference, we reoptimize this allocation on the topology found under mean demand rather than searching again for a topology under fluctuations.

Candidate graphs are Delaunay triangulations of points sampled uniformly in a unit disk, with the source at its center. For each of the sizes *N* = 400 and 700, the main campaign uses domain seeds 3, 4, and 5. The tree search begins from eight perturbed shortest-path trees, then alternates edge reweighting by marginal concave flow cost with edge exchange. Every steady-demand ratio in the numerical summary is measured against the best tree recorded for the same *b*, size, and domain. Although the search reaches that reference in all 30 combinations, it remains a numerical benchmark rather than a certificate of global optimality.

### Appendix B: Protocols, convergence, and numerical sensitivity

Parts of the simulation and analysis code implementing Eqs. (A1)–(B1), including the tree-search and adaptation routines, were drafted with assistance from Claude Opus 5 (Anthropic) and subsequently checked by the authors. The campaign uses *b*∈ {1, 3*/*4, 2*/*3, 1*/*2, 1*/*4} ; 2*/*3 is stored as 0.666667. Each explicit update is

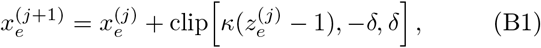

where *j* labels the iteration, clip bounds the increment, *κ* = 0.2, and *δ* = 0.02. An edge is removed once *r*_*e*_ *<* 10^*−*3^ max_*j*_ *r*_*j*_. Convergence requires max_*e*_ |*z*_*e*_ −1| *<* 10^*−*6^ under steady loading and 10^*−*5^ under fluctuating loading; these thresholds are stopping criteria, not error bounds on the continuous trajectory.

Peripheral growth adds sinks in 3, 6, 12, or 24 stages, ordered by distance from the source. The regrowth protocol reseeds every admissible edge with zero radius, whereas the control seeds an edge only once. At the first stage, seeded radii are *r*_seed_*/*2 with *r*_seed_ = 0.05 by default; later stages use half the median positive radius. We also run a sensitivity series with *r*_seed_ = 0.25. In the isotropic protocol all sinks are present from the start, while lengths scale from 0.2 to 1 over 6 or 12 stages. Symmetry of the static problem does not guarantee that these discrete histories select the same topology.

The summary contains 30 searches, 30 shear checks, 30 dense adaptations, 240 random-start adaptations, 420 growth runs, and 360 fluctuating-demand runs. All 1050 adaptation records converge, as does every scheduled stage in the growth records. The fluctuating-demand series uses *σ*∈ {0.5, 1, 1.5, 2, 3, 4} and contains 180 matched pairs with and without budget adjustment. Budget-adjusted runs must also reach a relative budget error below 2 × 10^*−*3^; the largest recorded error is 2.65 × 10^*−*6^.

Means in the text are computed separately for each network size across three domains. The random-start means additionally include eight initializations per domain, which should not be treated as independent spatial domains. Reported spreads describe variation between domains rather than confidence intervals. Figure 2(f) pools both sizes, and its loop-count error bars are sample standard deviations across the three domains.

For completeness, the dense-start mean excesses for *b* = 1, 3*/*4, 2*/*3, 1*/*2, 1*/*4 are 22.4, 35.7, 43.4, 59.0, 99.1% at *N* = 400 and 26.3, 48.9, 60.4, 83.0, 144.3% at *N* = 700. Random-start values range from 38.2 to 159.8% and from 38.3 to 178.8%, respectively. For *N* = 700 and *σ* = 2, the mean loop counts are 47.7, 25.7, 19.0, 9.3, 0 in descending order of *b*; at *σ* = 4, they are 87.0, 66.3, 57.3, 46.0, 26.3.

The main campaign uses *κ* = 0.2 and a step cap *δ* = 0.02. To test the central comparison at *b* = 1, we repeated the dense-start and 24-stage growth protocols with a fourfold smaller cap, *δ* = 0.005, while holding *κ* and the stopping criterion fixed. The dense excess changes from 24.6% to 18.3%; the staged excess moves from 8.5% to 7.2%. The coarse step shifts the absolute values but not their ranking, leaving staged growth much closer to the reference. Since the same ordering appears for all five values of *b* in the full campaign, this refinement supports the qualitative comparison on which our conclusion rests. More generally, the change of variable *t*^*′*^ = *κt* removes a constant *κ* from Eq. (6); any dependence of a numerical endpoint on *κ* must come from discretization and pruning rather than the continuous dynamics.

### Appendix C: Descent identity

For fixed lengths, 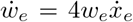. Equation (3) implies 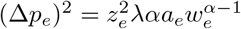, so

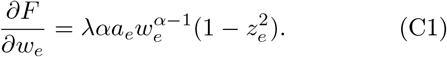

Multiplying Eq. (C1) by 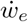 and summing gives Eq. (7), because 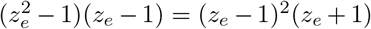 and *α* = *b/*2. The same argument applies to fixed *Q* with root-mean-square shear. The derivation assumes positive conductances on an admissible support and does not apply at the moments when that support changes.

### Appendix D: Estimating the maintenance exponent

Equation (3) constrains the mean shear profile through the product (1 −*b*)(1 +*q/*2), so a measured small-to-large shear ratio cannot separate *b* from the length–radius exponent *q*. Residual variation around the profile supplies an independent constraint: after the radius dependence is removed, the shear residual is proportional to the length residual, with a factor set by *b*. Over the range 0 *< b* ≤1 studied here, their standard-deviation ratio identifies *b* through Eq. (D3); the geometry-induced residual dispersion vanishes at *b* = 1.

Choose fixed reference units for shear, radius, and length, and fit

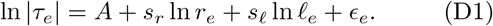

Here *A* is an intercept, *s*_*r*_ and *s*_*l*_ are fitted slopes, and *ϵ*_*e*_ is the residual. Equation (3) predicts *s*_*r*_ = *b* − 1 and *s*_*l*_ = (*b* − 1)*/*2, giving two estimates,

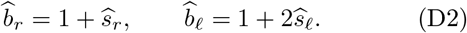

Agreement between the two estimates tests the model, although their covariance matters because radius and length may be strongly correlated. A separate geometric fit, ln *l*_*e*_ = *A*_*l*_ + *q* ln *r*_*e*_ + *η*_*e*_, defines *q* through intercept *A*_*l*_ and residual *η*_*e*_; regression on radius alone would mix the two effects. For domain 3 at *N* = 400, the marginal slopes are − 0.310 and − 0.415 for *b* = 3*/*4 and 2*/*3, respectively, compared with − 0.296 and − 0.406 at *N* = 700. Joint slopes recover the prescribed exponents to numerical precision.

For an exact model tree, let *ξ*_*e*_ be the residual of ln *l*_*e*_ after a free linear regression on ln *r*_*e*_, and let *ζ*_*e*_ be the corresponding residual of ln |*τ*_*e*_|. If sd(*ξ*) *>* 0, then

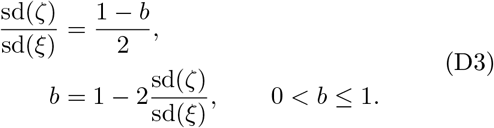

Here sd denotes standard deviation. The signed slopes in Eq. (D2) yield the same estimate and test the model more fully, whereas Eq. (D3) offers a complementary route independent of the mean length–radius trend. Using the unadjusted spread of ln *l* would overestimate the pre-dicted dispersion. An empirical test would require segment length, radius, and either flow or pressure drop to be measured in the same vessels, with pulsatility, viscosity variation, and measurement uncertainty taken into account.

